# Bio-sintering of Limestone to Produce Pollution Free Cement

**DOI:** 10.1101/2025.05.19.654766

**Authors:** Raja Murugan, Anant Dubey, Navdeep K Dhami, Abhijit Mukherjee

## Abstract

Industrial cement is produced by consuming too much energy and emitting very high CO_2_. Nature too produces cement, but it is pollution-free. This research demonstrates an emulation of natural cement by bio-sintering limestone, a process of successive dissolution and reprecipitation of limestone with the help of bacteria in ambient environmental conditions. When the bacteria *Acetobacter aceti* were introduced into a mixture of ethanol and limestone powder, the pH came down rapidly, and dissolution of limestone into calcium acetate was observed. After the ethanol was fully consumed, the pH rose gradually, and biocement crystals were reprecipitated. All the reaction rates have been determined, and the products have been characterised. It is observed that the morphology of the biocement could be tailored by controlling the reprecipitation process. The dissolution and reprecipitation took place within the time scale of present construction methods. Thus, this process can potentially manufacture industrial cement from limestone sans the pollution.

## Introduction

Concrete is an essential material for building roads, bridges, schools, and hospitals that sustain and improve the quality of human life, even in the remotest locations of the world. However, CO_2_ emissions from the annual consumption of 30B tonnes of concrete are causing nearly 8% of global CO_2_ emissions and threatening sustainability of the planet^1^. Cement production and consumption (see Fig. 1a) can broadly be divided into three phases: 1) *decomposition* of limestone (CaCO_3_) to create cement; 2) *transportation* of cement to the construction site; and 3) *cementation* to resolidify cement into concrete. Decomposition of limestone produces nearly 70% of the CO_2_ emission ^2^. It is performed in a kiln at around 1450 °C, and each molecule of CaCO_3_ generates a molecule of CO_2_, which is released to the atmosphere. Near-zero emissions from cement are essential for net-zero construction.

**Fig. 1.**
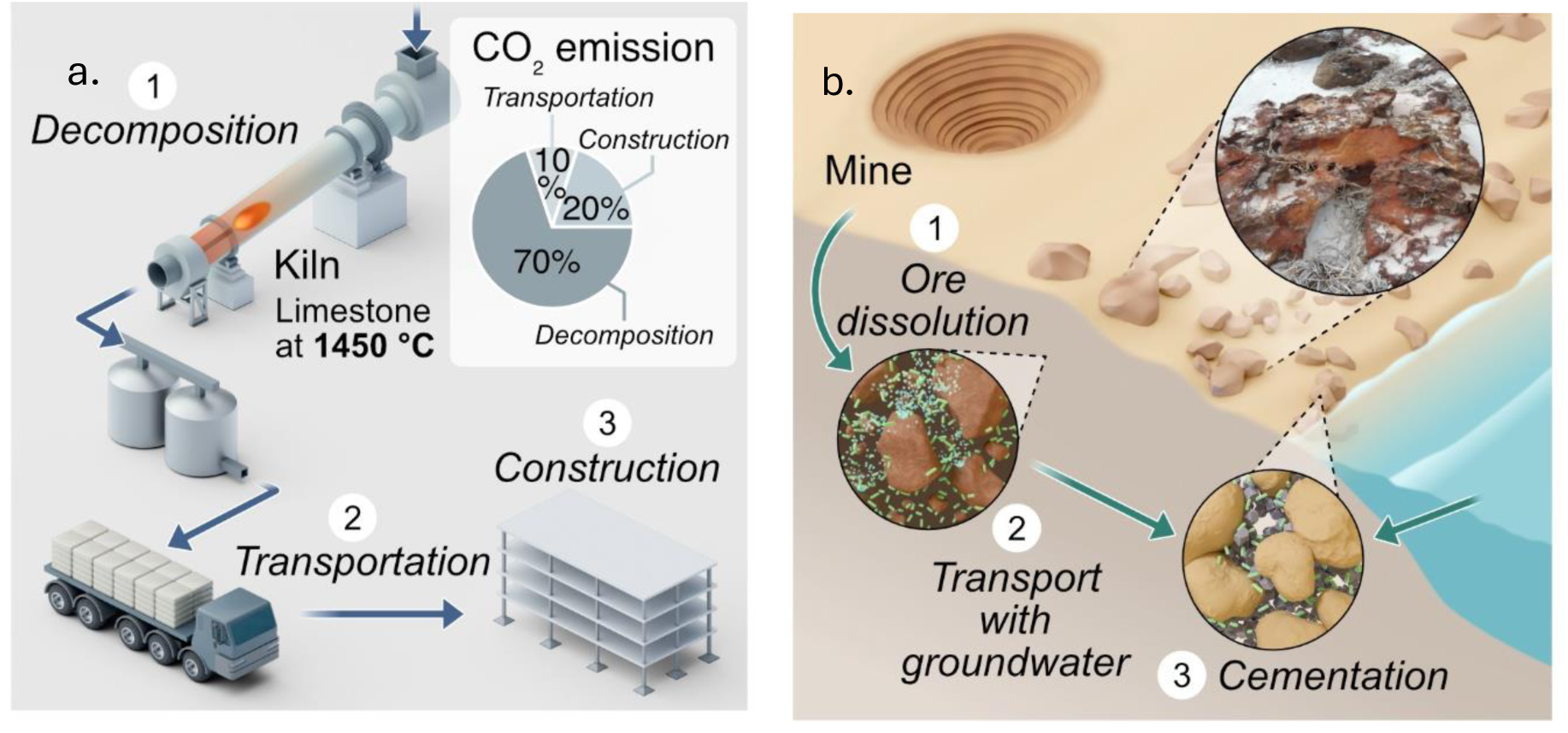
Building practices with limestone, a: Industrial; b: Natural.

Nature uses an environmentally benign method of building with CaCO_3_ in formations such as corals, stromatolites, stalagmites, and stalactites^3^. The role of bacterial mineralization in such formations was reported in 1996^4^. Ehrlich^5^ noted that mineral formation or dissolution is exploited by microbes if it favours their survival. Engineers found applications of this phenomenon in the fracture remediation of rocks^6^ and in remedying concrete^7^. Since then, a plethora of applications of the process have been reported in soil strengthening^8^, bio-concrete^9^, and self-healing concrete^10^. However, it is restricted to only the *cementation* phase, where commercial calcium and carbon sources have been used as raw materials. Our prior study shows that the raw materials, such as calcium chloride and urea, have substantial carbon footprints^11^. Moreover, this process produces harmful byproducts such as ammonia. To achieve the full benefit of natural cementation, unprocessed natural input material must be used. There is no report hitherto of a fully biological cement produced from limestone. However, evidence of the biodegradation of limestone in rocks, sediments, caves, and geological formations is available^12^. Successive decomposition and recombination in the formation of sponge spicules have been observed, which has been named bio-sintering^13^. We have observed the natural phenomenon prevalent in the beach rocks of Western Australia^14^, where bacteria cause the dissolution of the minerals at the mine site; the product is transported through the soil and then mineralised at the beaches to form beach rocks (see Fig. 1b). This phenomenon gives us a clue for bio-sintering of limestone. Hitherto, there is no report of bio-sintering with a view to developing an alternative to industrial cement. The challenge of emulating the natural bio-sintering into construction is that the new process must fit into the time scales of industrial cement manufacturing and building with concrete.

This investigation is the first attempt to bio-sinter limestone to produce cement directly from limestone following the acetate pathway. This pathway is a cleaner and environmentally benign alternative to the popular method of biocement production. Biocement is produced most popularly following the ureolytic pathway that reacts calcium chloride (CaCl_2_) with urea, which has a considerable carbon footprint. We bio-sinter naturally abundant limestone which would drastically reduce the cost^15^ and emissions from biocement^16,17^. In addition, this pathway avoids the generation of environmentally hazardous ammonia gas produced in the ureolytic system. Moreover, the chloride ions from CaCl_2_ that can cause corrosion of steel rebars in concrete are absent in this system^18^.

Limestone was decomposed in four days and reprecipitated in ten days. These are in line with the present time requirements of industrial cement making and concrete casting. Additionally, our investigation indicates that the biocement produced through this process is amenable to control of morphological and phase characteristics. Thus, it is possible to tailor the properties of the cement.

### The Proposed Bio-sintering

Bio-sintering of CaCO_3_ is a biological process where microorganisms induce the dissolution and reprecipitation of CaCO_3_ to form biocement. A series of biochemical reactions occurs in the bio-sintering mechanism via the complete oxidation of ethanol. Initially, *Acetobacter aceti* oxidizes ethanol (C_2_H_5_OH) in the presence of oxygen (O_2_), producing acetate ions (CH_3_COO⁻), protons (H⁺), and water (H_2_O) (Equation 1). The CaCO_3_ then dissolves in the acidic environment, releasing calcium ions (Ca^2+^) (Equation 2). The acetate ions are further oxidized by oxygen, forming additional CO_2_, water, and hydroxide ions (OH⁻) (Equation 3). The hydroxide ions combine with CO_2_ to form carbonate ions (CO_3_^2-^) (Equation 4), which then react with calcium ions to reprecipitate CaCO_3_ (Equation 5). Thus, the overall bio-sintering reaction can be summarized as the oxidation of ethanol leading to the dissolution of CaCO_3_, followed by acetate oxidation, carbonate ion formation, and the reprecipitation of CaCO_3_.

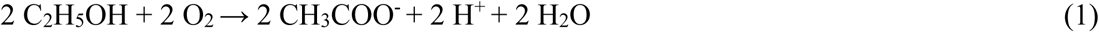

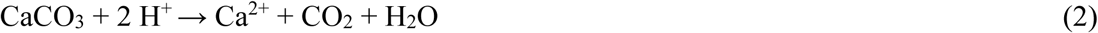

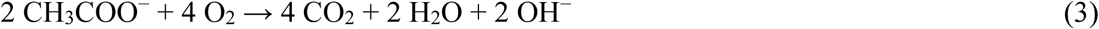

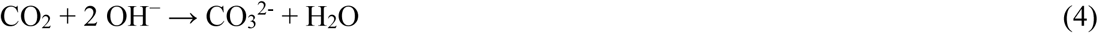

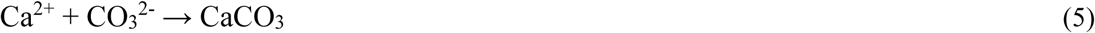

## Results

### Dissolution and Reprecipitation of CaCO_3_

The bio-sintering process using *Acetobacter aceti* (ATCC 15973) was investigated to evaluate the feasibility of microbially induced dissolution of limestone, followed by cementation of CaCO_3_ under ambient conditions. To decouple the microbial oxidation of ethanol from mineral interactions, an initial control experiment was performed without CaCO_3_ (Fig. 2a). In this system, a steady decline in ethanol concentration was observed, with 90% of ethanol oxidized in 60 hours. This conversion led to a corresponding increase in the concentration of acetate and a reduction in pH to 3.5, suggesting that microbial oxidation of ethanol produces sufficient acidity to potentially dissolve CaCO_3_.

**Fig. 2:**
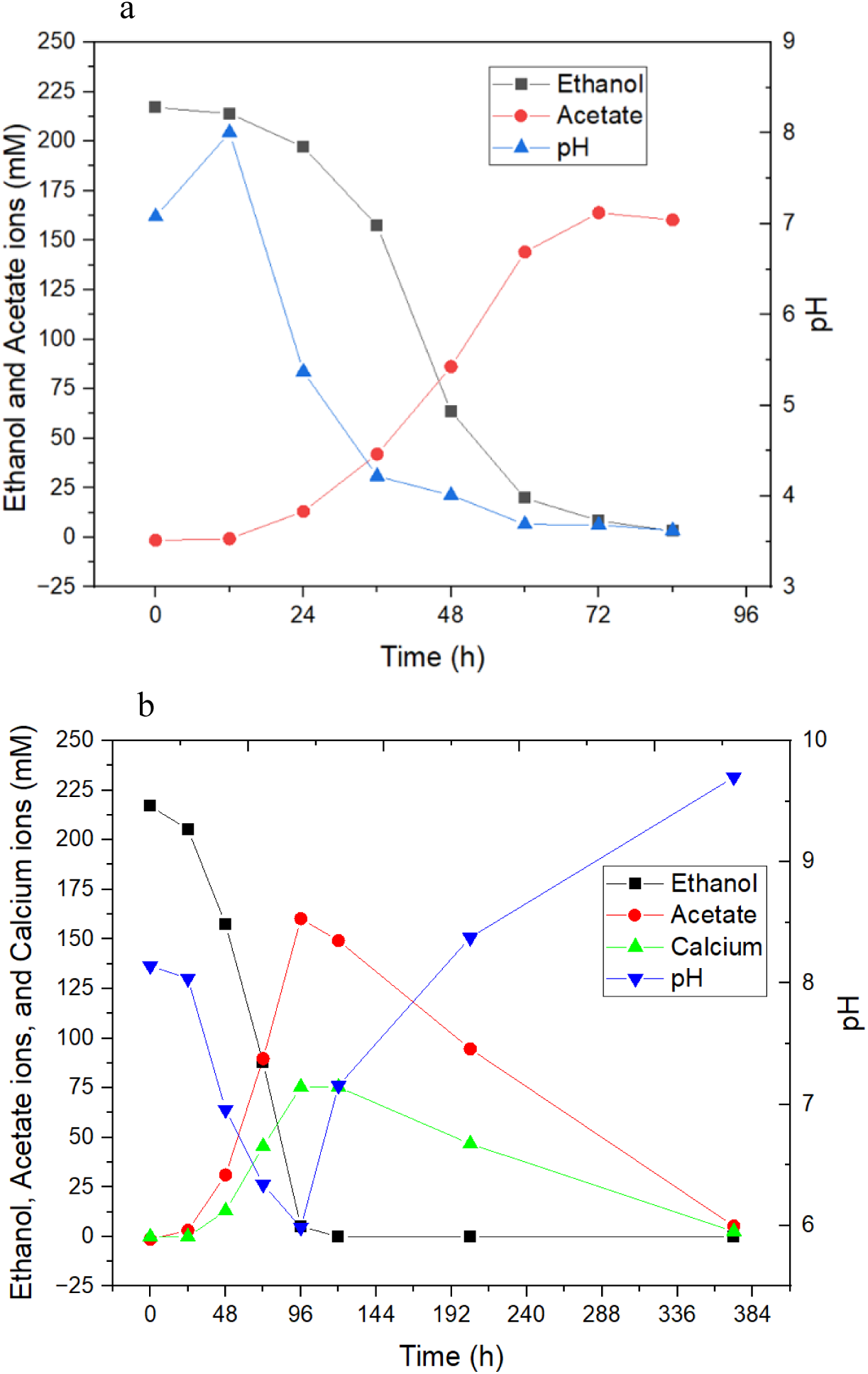
Bio-sintering of CaCO_3_ via ethanol and acetate oxidation using A. aceti. a) oxidation of ethanol to produce acetate. b) biological dissolution and reprecipitation of CaCO_3_ via ethanol and acetate oxidation, respectively.

Table 1 presents the chemical composition of limestone powder (Thermo Fisher Scientific) used in this experiment. 10 g/L of limestone powder (approximately 100 mM CaCO_3_) was included in the above bio-sintering medium to observe possible dissolution and reprecipitation. A series of contrasting biochemical transformations were observed, labelled as the Dissolution Phase and the Reprecipitation Phase (Fig. 2b). During the dissolution phase (0-96 h), ethanol oxidation by *A. aceti* progressed steadily, with undetectable ethanol concentration by the 96^th^ hour. Simultaneously, the concentration of the acetate rose above 150 mM, and the pH dropped to 5.96. These acidic conditions facilitated the dissolution of limestone powder, as evidenced by the rise in soluble calcium concentration to a peak of 76 mM. The calculated ethanol oxidation and acetate production rates were 2.87 mM/h and 2.33 mM/h, respectively, while the CaCO_3_ dissolution rate peaked at 1.23 ± 0.2 mM/h. This dissolution is consistent with the surface reaction between acetic acid and CaCO_3_, in line with previously reported microbial acidification-driven carbonate solubilization mechanisms^19,20^.

**Table 1.** Chemical composition of the limestone powder used for this study, determined by quantitative XRD analysis.

| Chemical Name | Chemical Formula | % |
| --- | --- | --- |
| Calcium carbonate | $\text{CaCO}_3$ | 94.4 |
| Calcium magnesium carbonate | $\text{Ca(Mg)CO}_3$ | 3.3 |
| Calcium silicate | $\text{Ca(Si)O}_3$ | 1.7 |
| Silicon dioxide | $\text{SiO}_2$ | 0.5 |
| Potassium magnesium aluminium silicate | $\text{KMg}_3\text{AlSi}_3\text{O}_{10}(\text{F,OH})_2$ | 0.1 |

Following the dissolution phase, the system transitioned into the Reprecipitation phase, marked by consumption of the acetate and an increase in the pH. From 96 to 392 hours, acetate was gradually oxidized, driving the pH up to 9.7 and reprecipitating CaCO_3_ back in the mineral form. This shift resulted in the complete removal of free calcium ions from the solution. The rate of acetate oxidation during this phase was 0.565 mM/h, while the CaCO_3_ precipitation rate was 0.23 ± 0.06 mM/h. These findings align with established principles of microbial-induced carbonate precipitation, where rising pH facilitates Ca^2+^ and CO_3_^2-^ recombination into solid CaCO_3_^21,22^

The central role of *A. aceti* in this bio-sintering process can be attributed to its well-characterized metabolic pathway. The bacterium utilizes membrane-bound pyrroloquinoline quinone (PQQ)-dependent alcohol dehydrogenase (ADH) and aldehyde dehydrogenase (ALDH) to oxidize ethanol to acetate. Once ethanol is exhausted, acetate is further metabolized via acetyl-CoA synthetase (ACS) into acetyl-CoA, feeding into the TCA cycle for energy generation^23^. This acetate oxidation not only provides metabolic energy but also contributes to alkalinization of the medium, a phenomenon previously associated with the microbial oxidation of negatively charged organic acids^20^.

Together, these results underscore the effectiveness of *A. aceti* in facilitating both dissolution and precipitation of CaCO_3_, offering a low-energy, biologically driven strategy for biocement production. The time taken for the dissolution of limestone (four days) and its reprecipitation (ten days) was compatible with the present industrial practices. The findings highlight the potential solution to the high emissions and energy consumption of industrial cement manufacturing and concrete making.

### Confirmation of reprecipitation of CaCO_3_ using FTIR

To assess the chemical composition of the biocement produced via bio-sintering, FTIR analysis was performed and compared against precursor limestone powder. As shown in Fig. 3, the FTIR spectra of both the microbially induced biocement and the reference limestone powder exhibit identical peaks at 1398, 871, and 712 cm^-1^. Similar peak assignments for CaCO_3_ have been reported in earlier studies^24,25^. These peaks correspond to the asymmetric stretching (ν₃) of the C–O bond, out-of-plane bending (ν₂), and in-plane bending (ν₄) modes of the carbonate (CO_3_^2-^) group, respectively. The close match in peak positions confirms that the mineral phase of the biocement is chemically consistent with that of precursor limestone powder.

**Fig. 3.**
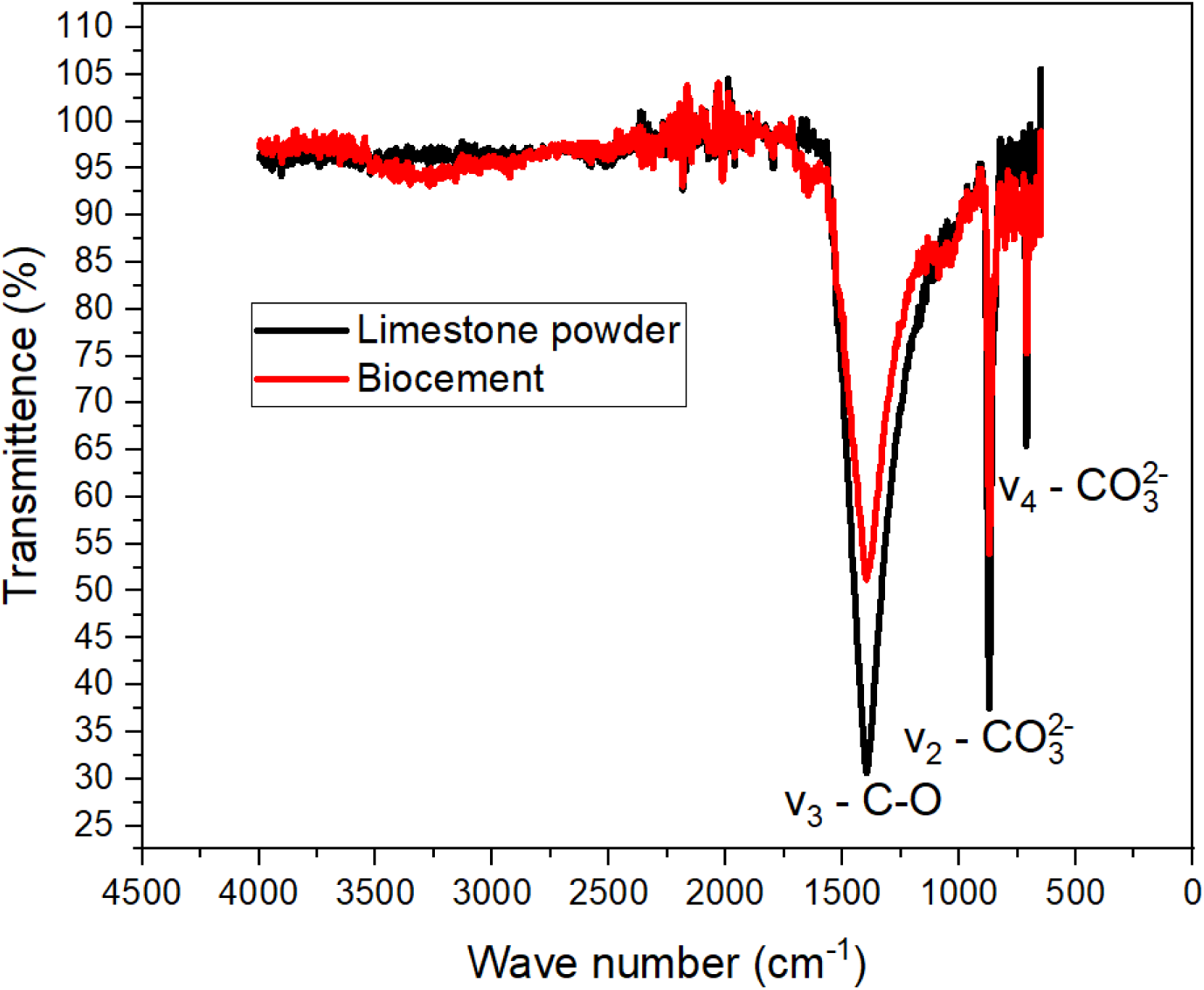
Fourier transform infrared spectroscopy (FTIR) spectrum of limestone powder and biocement produced via bio-sintering. Peaks such as 1398 cm^-1^ indicate asymmetric stretching (ν₃) of the C–O bond, and 871 and 712 cm^-1^ indicate out-of-plane bending (ν₂) and in-plane bending (ν₄) modes of the carbonate (CO_3_^2-^) group, respectively.

These findings indicate that the bio-sintering process leads to the formation of CaCO_3_ with a molecular structure and functional group profile equivalent to that of industrial-grade calcite. The reproducibility of characteristic peaks further supports the high purity and compositional fidelity of the biocement.

### SEM images illustrate the morphological evolution of CaCO_3_ crystals

Scanning Electron Microscopy (SEM) was used to investigate the morphological evolution of CaCO_3_ during the bio-sintering process. Fig. 4 presents representative SEM images captured at three stages: before dissolution, during dissolution, and after reprecipitation.

**Fig. 4:**
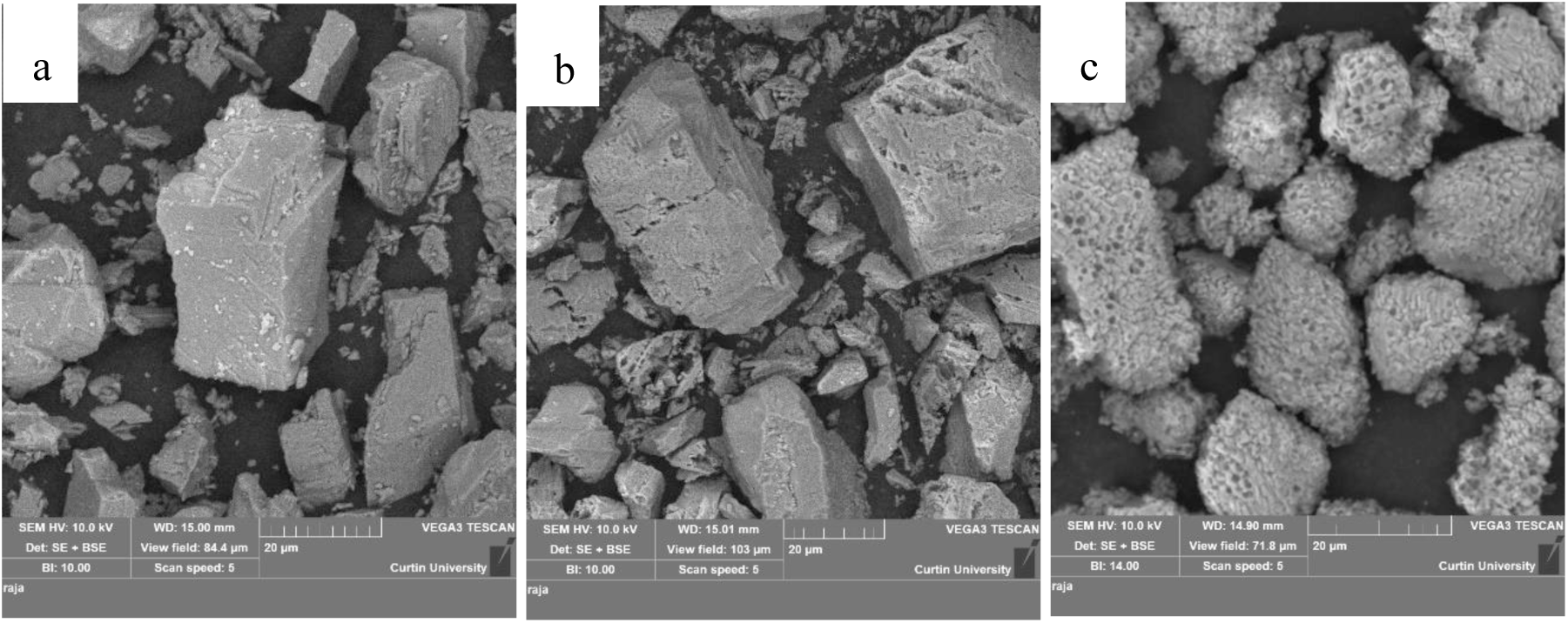
Scanning Electron Microscope (SEM) images of CaCO_3_. a) SEM image of CaCO_3_ before dissolution, b) SEM image of CaCO_3_ during dissolution, and c) SEM image of CaCO_3_ after reprecipitation (Biocement).

Before dissolution (Fig. 4a), the CaCO_3_ crystals exhibited well-defined rhombohedral morphology, characteristic of pure calcite, with smooth, pore-free surfaces. This morphology aligns with previously reported structures for synthetic or geological calcite^26^.

During the dissolution phase (Fig. 4b), substantial surface alterations were evident. The CaCO_3_ particles showed numerous pores and surface etching, attributed to the localized acidic microenvironment generated by the microbial oxidation of ethanol to acetic acid. This acid-mediated dissolution process led to the release of Ca^2+^ and CO_3_^2-^ ions, as reflected in the porous and degraded crystal structures. Such morphological changes enhance the surface area of the crystals and accessibility for microbial activity, facilitating accelerated dissolution.

After the reprecipitation phase (Fig. 4c), the CaCO_3_ crystals exhibited a transformation into aggregated, spheroidal to ellipsoidal forms with relatively rough, perforated textures. These sponge-like features are indicative of microbial mediation during the reprecipitation of CaCO_3_. The rise in pH due to acetate oxidation by *A. aceti* promoted supersaturation and subsequent crystal nucleation. The resulting morphology, marked by rounded, aggregated particles, is a well-documented outcome of biologically induced mineralization processes^27,28^.

The shift from rhombohedral to spheroidal crystal forms suggests that crystal growth dynamics can be tailored by bio-sintering. Factors such as microbial species, medium composition, saturation index, and the presence of ions like magnesium are known to influence crystal morphology during biocementation^29–31^. This aspect requires further investigation. The qualitative EDS spectra of the limestone powder and biocement are provided as supplementary material (Fig. S1). Both spectra confirm the presence of carbon, calcium, and oxygen, indicating that the primary constituent is CaCO_3_.

### XRD confirms that the reprecipitated biocement is calcite

Quantitative X-ray diffraction (XRD) analysis was performed to investigate the phase composition and crystallographic characteristics of the reprecipitated biocement. The analysis compared the diffraction patterns of biocement samples with those of precursor limestone powder (Fig. 5). The XRD spectra of the biocement showed a high degree of similarity to that of the limestone powder, with both displaying prominent peaks corresponding to the calcite phase. This observation confirms that the primary crystalline phase formed during reprecipitation is calcite, the most thermodynamically stable polymorph of calcium carbonate. Phase quantification indicated that the limestone powder consisted of approximately 95% calcite, with the remaining fraction identified as dolomite [Ca(Mg)(CO_3_)_2_]. Similarly, the biocement also comprised more than 95% calcite, with a minor fraction of dolomite detected.

**Fig. 5.**
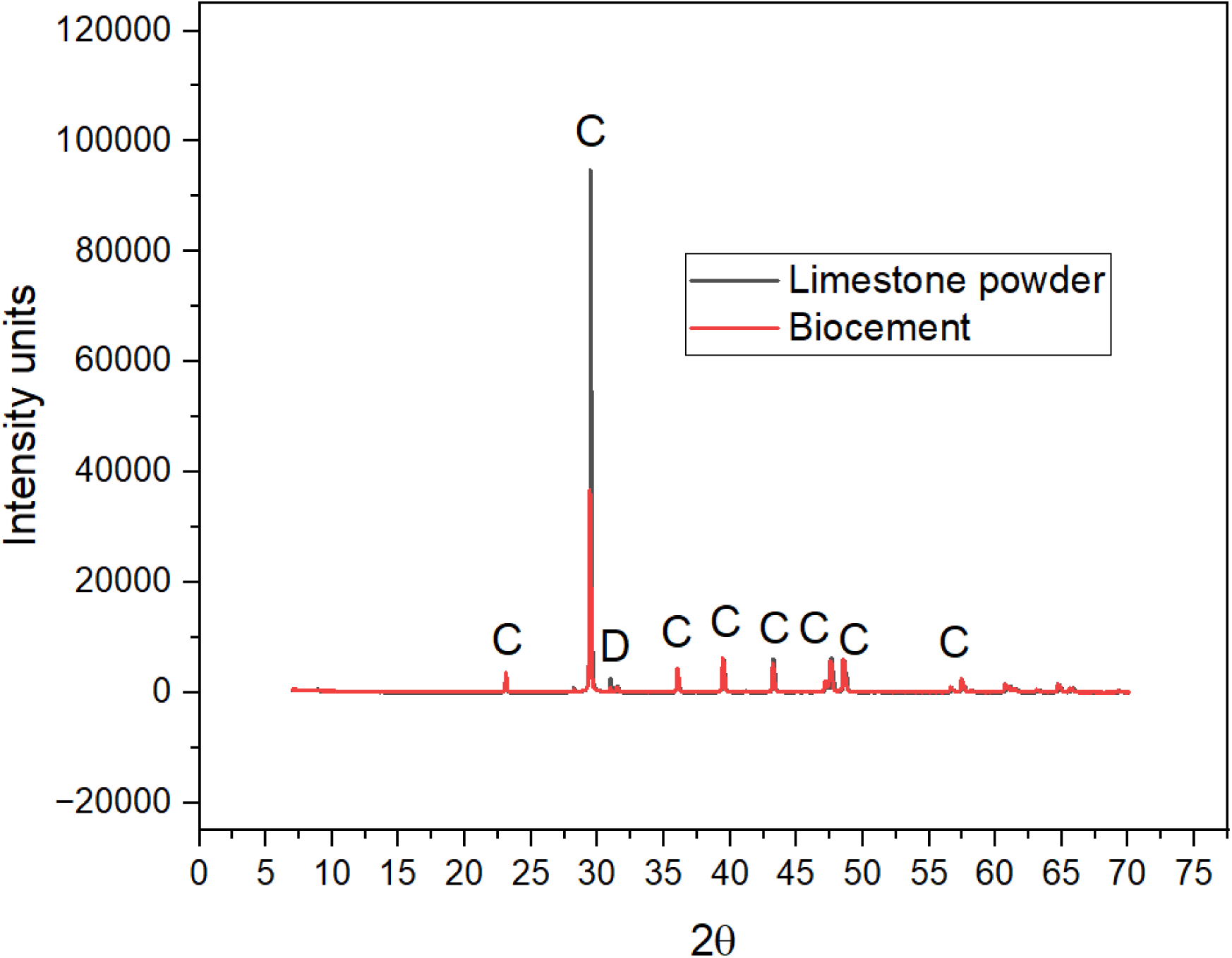
Quantitative X-ray diffraction (XRD) spectrum of limestone powder and biocement produced via bio-sintering process. C – Calcite and D – Dolomite.

The presence of dolomite in both samples may be attributed to trace amounts of magnesium ions present in the starting limestone powder used for the bio-sintering process. Magnesium incorporation into calcium carbonate structures is commonly observed in natural minerals and is known to influence crystal formation and stability^32^. The microbial species involved, along with the ionic composition of the medium, play a crucial role in determining the resulting CaCO_3_ polymorph. In this study, *Acetobacter aceti* facilitated the formation of calcite, aligning with previous reports that link microbial activity and solution chemistry to calcite precipitation^33^.

These results confirm that the bio-sintering process effectively produces biocement primarily composed of calcite with minor impurities, yielding a material that is structurally and chemically comparable to the precursor limestone powder. The high calcite content suggests the material is suitable for structural and environmental applications where stability and durability are essential^34,35^.

### Particle size analysis shows the size distribution of biocement

Particle size distribution measurements were carried out to evaluate the dimensional changes in CaCO_3_ particles induced by the bio-sintering process. Fig. 6 presents the comparative size distribution curves of limestone powder and the biocement formed after microbial treatment. The d(0.1), d(0.5), and d(0.9) values for the limestone powder were measured as 3.58 µm, 13.72 µm, and 31.29 µm, respectively, indicating a broad particle size distribution with relatively larger crystal aggregates.

**Fig. 6.**
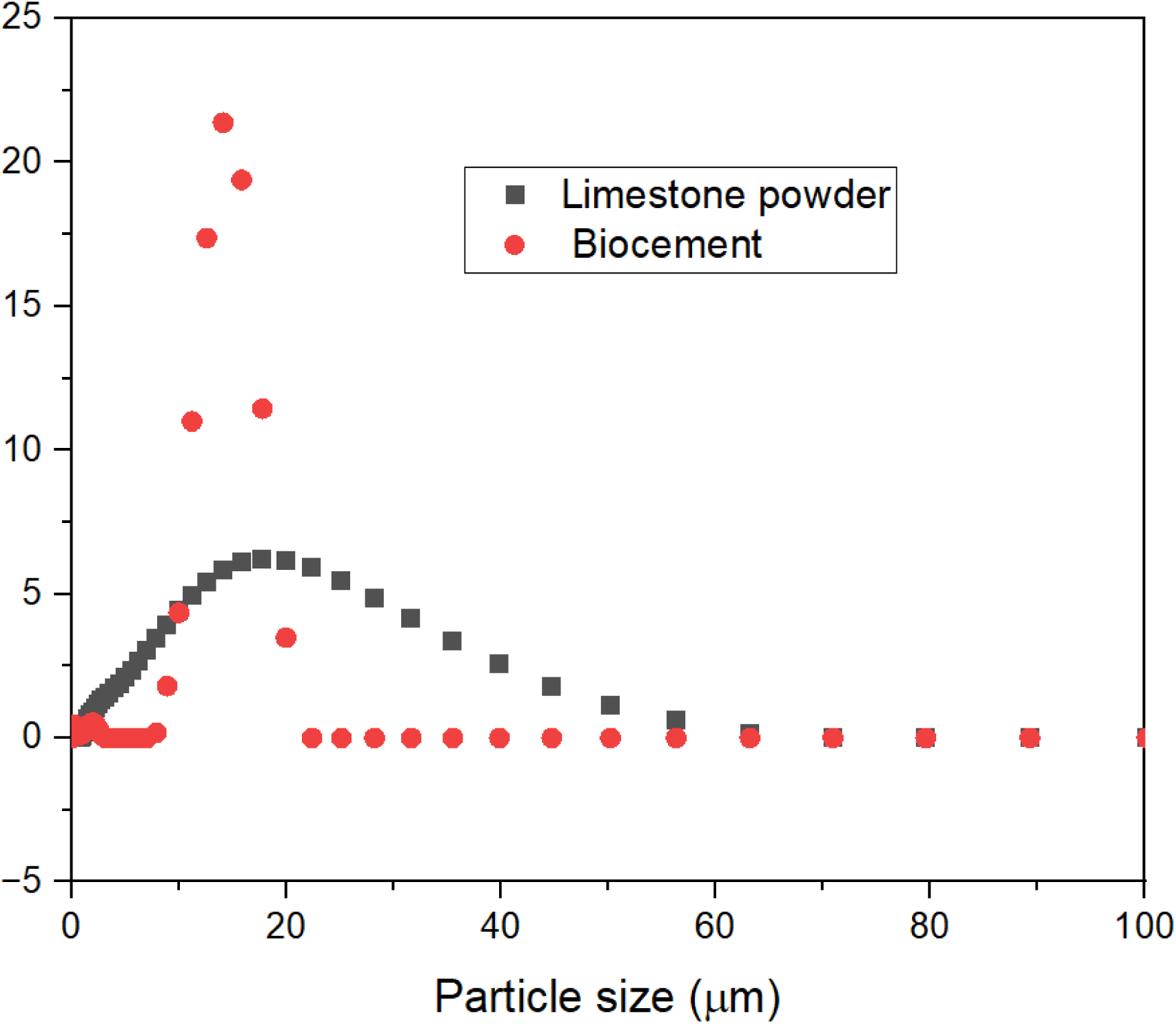
Particle size distribution curve of limestone powder and biocement produced via bio-sintering process.

In contrast, the biocement exhibited altered particle size characteristics, with d(0.1), d(0.5), and d(0.9) values of 8.13 µm, 13.02 µm, and 16.50 µm, respectively. This shift suggests a narrowing of the particle size distribution and the enhanced size uniformity after bio-sintering. The observed reduction in d(0.9) and increase in d(0.1) point toward homogenizing the particle population, which may result from the dissolution of the initial CaCO_3_ crystals followed by controlled microbial reprecipitation.

These changes are consistent with the thermodynamically driven nucleation and crystal growth mechanisms, wherein increased supersaturation during microbial activity can reduce the critical nucleus size, leading to the formation of smaller, more uniform crystals^36^. The resultant biocement morphology reflects the influence of microbial mediation on precipitation kinetics and crystal development. Further, by adjusting environmental conditions during the cementation process, it may be possible to influence the particle size, which in turn could help shape the physical properties of the final material.

In conclusion, this study demonstrates the successful production of biocement through the bio-sintering process, utilizing *Acetobacter aceti* for microbial dissolution and reprecipitation of CaCO_3_. The bio-sintering method offers a sustainable alternative to traditional cement production, significantly reducing the energy consumption and CO_2_ emissions associated with present industrial cement manufacturing. The biocement produced through this method exhibits similar morphological, phase, and chemical characteristics to limestone powder. Results indicate that it should be possible to tailor the properties of the biocement by controlling the particle size through environmental adjustments in the reprecipitation phase.

Bio-sintering also opens the possibility of cementing granular materials by utilising the naturally present native minerals in sand or soil without having to augment synthetic minerals. Such a system would closely emulate nature’s way of construction. Further research is planned in two directions: 1) to optimize the process parameters and improve the scalability; and 2) to explore applications of bio-sintering in soil stabilization, coastal protection, and infrastructure development.

## Methods

### Growth and bio-sintering medium

*Acetobacter aceti* (ATCC 15973) was obtained from the American Type Culture Collection and revived on autoclaved Mannitol-Agar medium containing 5 g/L yeast extract, 3 g/L peptone, 25 g/L mannitol, and 15 g/L agar. Before the bio-sintering experiments, the bacteria were subcultured in a liquid medium with 10 g/L yeast extract.

In a 250 mL flask containing 100 mL of bio-sintering medium consisting of 1% (w/v) ethanol, 10 g/L limestone powder (Thermo Fisher Scientific), and 1 g/L yeast extract. The yeast extract was autoclaved separately, while ethanol and limestone powder were added afterward. The overnight-grown *A. aceti* culture in 10 g/L yeast extract medium was centrifuged, and the supernatant was discarded. The resulting cell pellet was resuspended in the bio-sintering medium to achieve an initial optical density (OD) of 0.2.

### Bio-sintering Process Monitoring

The bio-sintering process was monitored over time following the resuspension of bacteria into the bio-sintering medium. At various time intervals during the process, 2 mL of culture medium was collected and centrifuged at 12000 rpm for 5 minutes. The supernatant was then used to measure the concentrations of ethanol, soluble calcium, acetate, and pH. The pellet was analyzed using Scanning Electron Microscopy with Energy Dispersive X-ray Spectroscopy (SEM-EDS), X-ray Diffraction (XRD), Fourier Transform Infrared Spectroscopy (FTIR), and a particle size analyzer. The experiments were performed thrice, independently, to ensure reproducibility.

**Table 2.** presents the experimental monitoring and analysis of the bio-sintering process.

| Experimental Flask | Factors monitored | Biocement characterisation |
| --- | --- | --- |
| Bio-sintering medium: 1% ethanol (W/V), 1 g/L yeast extract, and with or without 10 g/L limestone powder | Concentration of ethanol, acetate, and soluble calcium, and pH. | FTIR, SEM-EDS, XRD and Particle size analyser |

### Analytical Techniques: Methods to measure the concentration of ethanol, acetate ions, calcium ions, and pH

Ethanol concentration was determined using the Ethanol Assay Kit (Abcam, ab272531) following a colorimetric method. Samples were diluted to fall within the standard range of 0 to 170 mM. In a 96-well microplate, 66.66 μL of each sample or standard was mixed with an equal volume of Reagent A and incubated at room temperature for 8 minutes. To stop the reaction, 66.66 μL of the stopping reagent (Reagent B) was added, bringing the final volume to 200 μL. Absorbance was measured using a microplate reader, and ethanol concentration was calculated from the slope of the standard curve generated by plotting absorbance against known ethanol concentrations.

Acetate concentration was measured using a colorimetric assay with the Acetate Assay Kit (Abcam, ab204719). Samples were diluted to fall within the standard range of 0 to 1 mM, and standard solutions were prepared accordingly. In a 96-well microplate, 50 μL of each sample or standard was mixed with 50 μL of a reagent mix containing acetate assay buffer, acetate enzyme mix, ATP II/ATP, acetate substrate mix, and developer solution in a 21:1:1:1:1 ratio, making a total reaction volume of 100 μL. The mixture was incubated at room temperature for 40 minutes, protected from light, before measuring absorbance at 450 nm using a microplate reader. A standard curve was generated by plotting absorbance against concentration, and the sample acetate concentration was determined using the slope of the standard plot.

The concentration of soluble Ca²⁺ was determined using the complexometric titration method^37^. To begin, 40 μL of the supernatant was diluted to 10 mL by adding 9.56 mL of water and 400 μL of 1N NaOH to raise the pH. Subsequently, a few drops of 1% (w/v) hydroxy naphthol blue disodium salt solution were added as an indicator. The titration was carried out using a 1 mM EDTA disodium salt solution, and the endpoint was reached when the colour changed from pink to blue. For calibration, standard solutions of 0 to 2 mM CaCl₂ were prepared and titrated against the 1 mM EDTA solution. The endpoints of these standard solutions were recorded, and a graph was plotted showing the relationship between the CaCl₂ concentration and the volume of 1 mM EDTA required to reach the endpoint. The concentration of the unknown sample was then determined from the slope of the resulting plot. The pH of the supernatant solution was measured using a pH meter (Thermo Scientific, Orion Star A211).

### FTIR Analysis

Fourier-transform infrared (FTIR) spectroscopy was conducted to analyze the precipitate obtained after the bio-sintering process. For comparison, precursor limestone powder used in the bio-sintering process was also subjected to FTIR analysis. Spectra were recorded using an FT-IR-100 spectrometer (Perkin Elmer) over the wavenumber range of 4000–650 cm⁻¹. The resulting spectra were utilized for peak identification.

### SEM analysis

Morphological analysis and elemental composition of the dried powder were carried out using Scanning Electron Microscopy (SEM) and qualitative Energy Dispersive X-ray Spectroscopy (EDS). The analysis was conducted with a Variable Pressure Electron Microscope (VP-SEM, Zeiss, EVO 40-XVP, 2008). Samples were mounted on carbon-aluminium tape, and their surfaces were coated with carbon using a carbon evaporative coater (Creissington, 2080C, 2011). The imaging was performed at a beam intensity of 10 and a voltage of 10 kV, with a working distance of around 15 mm. Both secondary electron (SE) and backscattered electron (BSE) imaging were combined to capture the electron micrographs.

### **X-** ray Diffractometry analysis

The samples were resuspended in ethanol and deposited onto low-background sample holders. X-ray diffraction (XRD) data were collected using a Bruker D8 Advance diffractometer equipped with Ni-filtered Cu Kα radiation (40 kV, 40 mA). Measurements were performed over the range of 7–120° 2θ with a step size of 0.015°. Phase identification was conducted using Bruker EVA 5.2 software and the COD (Crystallography Open Database). Phase quantification was performed using the Rietveld refinement method implemented in Topas Academic 7, with crystal structure data retrieved from the COD database

### Particle Size Analysis

The size distribution of CaCO₃ particles was measured using a Mastersizer (2000S). An appropriate amount of the sample was dispersed in deionized water, which was used as the medium. Measurements were conducted using laser diffraction, and the particle size distribution was recorded. The data were analyzed to obtain D10, D50, and D90 values.

## Acknowledgement

The authors acknowledge the financial support received from the Asian Office of Aerospace R&D through award number FA2386-24-1-4001. The characterisation of materials was performed at Curtin University’s John de Laeter Research Centre, and the synthesis of materials was performed at the Bio-Activated Materials Laboratory of Curtin University. The authors acknowledge the suggestions received from Dr Jason Folley, Lead, Biotechnology Advanced Development Team, AFRL Materials & Manufacturing Directorate (AFRL/RXEB), Wright-Patterson AFB, OH

## Author Contributions

R.M. designed and performed the experiments, analyzed the data, and wrote the manuscript. A.M. conceived the project, contributed to the design of the experiments, supervised the project, and wrote the manuscript. N.D. and A.D. each contributed to the conceptualization and overall development of the project.

## Material and Correspondence

Correspondence to **Abhijit Mukherjee**.

## Supplementary Details

Qualitative EDS spectrum of limestone and biocement (Fig. S1).

## Data availability

The data that support the findings of this study are available from the corresponding author upon reasonable request

