## Supplementary File S1 for "Bio-sintering of Limestone to Produce Pollution Free Cement"

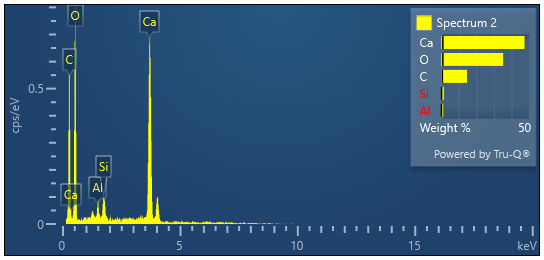
**Fig. S1 Qualitative EDS spectrum of limestone powder and Biocement**

Qualitative EDS spectrum of limestone powder

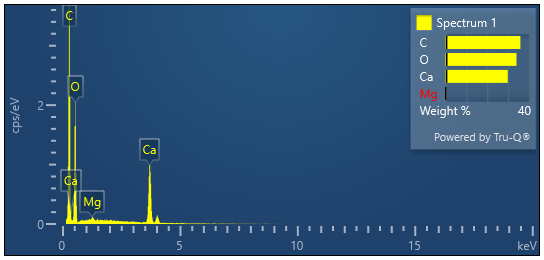

Qualitative EDS spectrum of biocement
